# Functional domain annotation by structural similarity

**DOI:** 10.1101/2023.01.18.524644

**Authors:** Poorya Mirzavand Borujeni, Reza Salavati

## Abstract

Traditional automated in *silico* functional annotation uses tools like Pfam that rely on sequence similarities for domain annotation. However, structural conservation often exceeds sequence conservation, suggesting an untapped potential for improved annotation through structural similarity. This approach was previously overlooked before the AlphaFold2 introduction due to the need for more high-quality protein structures. Leveraging structural information especially holds significant promise to enhance accurate annotation in diverse proteins across phylogenetic distances.

In our study, we evaluated the feasibility of annotating Pfam domains based on structural similarity. To this end, we created a database from segmented full-length protein structures at their domain boundaries, representing the structure of Pfam seeds. We used *Trypanosoma brucei*, a phylogenetically distant protozoan parasite as our model organism. Its structome was aligned with our database using Foldseek, the ultra-fast structural alignment tool, and the top non-overlapping hits were annotated as domains. Our method identified over 400 new domains in the T. *brucei* proteome, surpassing the benchmark set by sequence-based tools, Pfam and Pfam-N, with some predictions validated manually. We have also addressed limitations and suggested avenues for further enhancing structure-based domain annotation.

## 1 Introduction

Functional annotation is a critical process that seeks to delineate the identity of a protein in three distinct dimensions: its final location, its function, and the biological processes in which it participates. Experimental functional annotation can be time-consuming and expensive, underscoring the significant importance of automated *in-silico* functional annotations. These tools offer the convenience of high-throughput analyses without the need for manual inspection. Currently, most automated *in silico* annotation tools rely on sequence similarity to manually annotated proteins. Among them, annotations based on 1:1 orthology and InterPro2GO are particularly prevalent (Gene Ontology Consortium website). In 1:1 orthology-based annotation, sequence search tools like BLASTP are primarily used to infer orthology relationships. InterPro2GO, the most widely adopted automated annotation tool (Gene Ontology Consortium website), identifies different sequence signatures in the query protein using InterProScan (Paysan-Lafosse, Blum et al. 2022). The identified sequence signatures will be translated into GO terms using the InterPro2GO table, a curated table showing the associations between sequence signatures and the functional annotation of the proteins.

On the other hand, protein structure is shown to be three to ten times more conserved than its sequence (Illergård, Ardell et al. 2009). Consequently, it is anticipated that a larger number of proteins could potentially be annotated by integrating protein structure into the annotation process. Some studies have indeed succeeded in using protein structures to annotate domains of proteins that could not be annotated by sequence alone (Zarembinski, Hung et al. 1998, Li, Huang et al. 2018, Bartas, Slychko et al. 2022). However, these structure-based approaches have not been broadly adopted. One reason is the experimental resolution of protein structures has predominantly focused on well-studied proteins. Additionally, previous structure prediction tools were not sufficiently accurate, which made the prediction of unannotated protein structures challenging. Despite these limitations, these prediction tools have demonstrated potential for contributing to functional annotation (Zhang, Freddolino et al. 2017, Gligorijević, Renfrew et al. 2021).

Until recently, the structure-based functional annotation has not been the focal point for large-scale annotations mainly since only a limited number of protein structures had been resolved experimentally, and computational predictions were not sufficiently accurate. The introduction of Alphafold2 sparked a revolution in predicting protein structure. Alphafold2 is capable of predicting protein structures with an accuracy similar to experimental methods (Jumper, Evans et al. 2021, Varadi, Anyango et al. 2022), thus opening new avenues in the field of functional annotation.

With protein structures predicted, the next step is to identify proteins with similar structures. Although advanced tools like DALI (Holm 2022) and TM-align (Zhang and Skolnick 2005) are available for this purpose, they can be computationally intensive when used to search comprehensive structure databases. RUPEE (Ayoub and Lee 2019) is a faster tool, but it only looks for structural similarity and not sequence similarity. Foldseek, a newly developed structure search tool, can find structurally similar proteins significantly faster than other tools (van Kempen, Kim et al. 2023).

The combination of Alphafold2 and Foldseek may facilitate the annotation of a greater number of proteins. A recent study demonstrated that in the proteomic comparisons of species that are evolutionarily distant, there can be instances where the best reciprocal matches cannot be identified using sequence similarity methods. However, these matches can be detected when top reciprocal structural correspondences are considered (Monzon, Paysan-Lafosse et al. 2022). In a separate study, Ruperti et al. employed Foldseek to identify the closest structural match in model organisms to the query proteins and transferred the annotations from the top hits (Ruperti, Papadopoulos et al. 2023). Their findings were remarkable, as the use of Foldseek instead of sequence alignment tools could lead to the annotation of up to 50% more genes.

Bordin et. al also used Foldseek and SSAP to annotate the CATHe domains of 21 model organisms and identify new structural domains in them (Bordin, Sillitoe et al. 2023).

As previously mentioned, InterPro2GO uses sequence-based domain annotation by utilizing multiple member databases within InterPro. Each database holds a distinct type of sequence signature. For example, Pfam, one of the most widely used members, holds the sequence signature of protein functional domains (Mistry, Chuguransky et al. 2021). These sequence signatures are extracted from multiple sequence alignments of well-recognized domain instances known as Pfam seeds. Pfam employs Hidden Markov Models (HMM) to model these signatures, which can then be searched against proteins of interest using the hmmscan module of HMMER (Potter, Luciani et al. 2018). It is worth mentioning that Pfam also post-processes the hits and its post-processing might change the importance order of the hits reported by HMMER.

There are also tools for improving the sensitivity of Pfam annotation. For instance, by creating a profile of query proteins and aligning it with Pfam profiles, more hits can be identified, although the order of these hits might not necessarily reflect their relevance (Söding, Biegert et al. 2005). A recent study found that by employing Convolutional Neural Networks (CNNs) to store sequence signatures, over 9.5% more new domains could be annotated (Bileschi, Belanger et al. 2022). These new annotations are referred to as Pfam-N annotations.

In this study, we aim to explore a novel approach for Pfam domain annotation relying on Protein structure. As Pfam seeds represent a portion of a protein sequence, we created a database of Pfam seeds by dividing the corresponding full-length structures at their domain boundaries. From now on, we will refer to the source protein as the Full-Length Structure of Pfam Seed Source (FLSPSS). Next, we evaluated the reliability of the structural alignment by aligning the FLPSS with the constructed domain structure database to determine the frequency of structural alignment between different Pfam seeds.

We focused on *Trypanosoma brucei*, an early-diverged eukaryotic parasite, and aligned its structome with our domain database as a case organism. We benchmarked the predicted domains by comparing them to the Pfam v35.0 and Pfam-N predictions as the gold standard. Additionally, we conducted a manual review of some of the domains that were predicted for the proteins involved in *T. brucei’s* mitochondrial RNA editing and served as a case study.

## 2 Materials and Methods

### 2.1 Construction of a Pfam domains structure database (PfamSDB)

The FLPSSs of Pfam v35.0 were retrieved from the Alphafold database (AlphaFold DB, version 4). AlphaFold and Pfam use different versions of UniProt. As a result, for fewer than 0.2% of Pfam seeds, the sequence obtained by trimming the FLPSS based on the Pfam seed coordinates did not match the Pfam seed sequence. These instances were subsequently removed from the database. In the end, 6.4% of Pfam seeds were not present in the database, either because AlphaFold had not predicted their structure (6.2% of instances) or due to version discrepancies in the source sequence between Pfam and AlphaFold DB (0.2% of instances).

The FLPSSs were cut on their Pfam borders and stored the chopped domains in PDB format. For this task, the source codes of the PDB-tools package (Rodrigues, Teixeira et al. 2018) were modified to do cutting and format conversion to PDB simultaneously. GNU parallel was extensively used for parallelizing the computations (Tange 2018). PfamSDB contained over 1.1 million structures.

### 2.2 Benchmarking Foldseek probability

The FLPSSs were aligned with the PfamSDB using Foldseek v7-04e0ec8. The number of sequences passing the pre-filtering step was set to a high number by the “—max-seqs 1e9” option to get all possible alignments. A match with the same Pfam domain as the target seed instance was deemed a True Positive (TP), while a match with a different Pfam domain was considered a False Positive (FP). In our scoring system, for a result to be reported as a TP, the seed region on the query must be covered by more than 25% in alignment with an instance of the same Pfam domain. Conversely, if an instance of a different Pfam domain covers more than 25% of the seed region on the query in the alignment, it is considered an FP.

### 2.3 Annotating *T. brucei* structome and benchmarking against Pfam and Pfam-N

The structome of *T. brucei* was aligned with PfamSDB using the same parameters as the former step and specified the “--greedy-best-hits” option to report non-overlapping highest-scoring hits. Our labeling approach was similar to the one employed previously, with one key difference: in this phase, we used regions annotated with either Pfam or Pfam-N as our gold standard for comparison, instead of using the seed regions that served as the gold standard in the previous step. If the domain predicted by the gold standard was not retrieved, it was labeled as False Negative (FN). We used Seaborn (Waskom 2021) and matplotlib (Hunter 2007) for visualization.

### 2.4 Pfam domain annotation by MMseqs2

To investigate the added value of structural similarity for domain annotation, the same procedure as explained above was done by searching the sequences of *T. brucei* proteome against sequences of PfamSDB by MMseqs2 v14-7e284 (Steinegger and Söding 2017), the same program used by Foldseek under the hood. The “-s 8.5” parameter was specified to run it with high sensitivity as Foldseek uses MMseqs2 in high sensitivity mode.

### 2.5 Pfam domain annotation by HMMER

As mentioned earlier, Pfam annotation relies on the post-processing of HMMER hits. Postprocessing relies on some curated data such as the score threshold for each Pfam domain. As we did not have such data for Foldseek alignment scores, we also evaluated Pfam domain annotation by simply selecting the best-ranking domains reported by HMMER. In this regard, we removed the seed instances that were missing in the PfamSDB and made the HMM database using the “hmmbuild” module. The “hmmscan” was used for aligning the *T. brucei* proteome with the adjusted Pfam database. Hits with e-values up to 0.001 were considered for the rest of the analysis.

## 3 Results and discussions

### 3.1 PfamSDB Contains High-Confidence Short Structures

AlphaFold reports the estimated confidence level for each residue as the predicted Local Distance Difference Test (pLDDT). The pLDDT value ranges between 0 and 100, and higher values indicate more confident predictions. Early observations have shown that there is a high overlap between residues with low pLDDT and regions known as Intrinsically Disordered Regions (IDRs) that do not fold into specific structures (Jumper, Evans et al. 2021). According to the AlphaFold website, residues with a pLDDT above 90 are expected to be highly accurate, while regions with a pLDDT between 70 and 90 are considered to have good backbone prediction. If low pLDDT regions correspond to IDRs, we do not expect accurate structural matches for those regions. Furthermore, the significance level of structural alignment hits has been shown to depend on protein length. Monzon et al. have demonstrated that Foldseek has difficulty establishing relationships for some short proteins (those with fewer than 200 amino acids) when identifying the best reciprocal hits (Monzon, Paysan-Lafosse et al. 2022).In contrast, sequence-based aligners do not exhibit this problem.

As mentioned, pLDDT is reported per residue, however, for the purposes of this discussion, we will refer to the average pLDDT of a region as avg_pLDDT. Figure 1A depicts the average of the avg_pLDDT values of instances of each Pfam domain, while Figure 1B illustrates the distribution of the average size of instances of each Pfam domain. Overall, the instances exhibit a high avg_pLDDT, and the average length of instances across different Pfam domains typically falls below 200. To be more specific, the third quartile (75^th^ percentile) for the average size of instances for each Pfam is 209.

**Figure 1.**
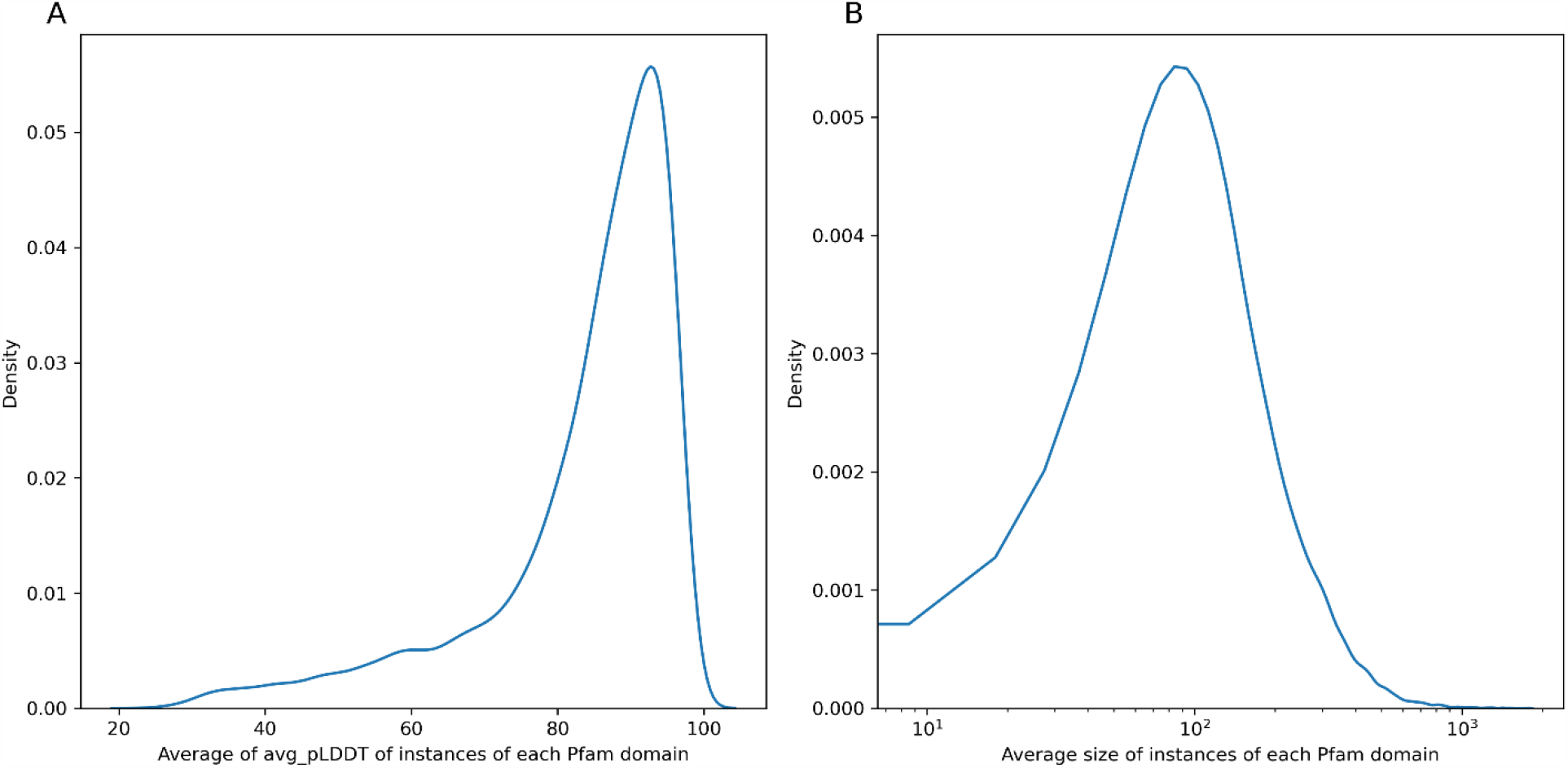
The kernel density estimation of the average of avg_pLDDT (A) and average of size of instances (B) of each Pfam domain.

### 3.2 TPs and FPs can be separated based on bitscore

We aimed to assess the frequency with which instances of one Pfam domain align to instances of a different Pfam domain, identified as FPs. We then sought to determine if Foldseek probability could distinguish these FPs from TPs. Using Pfam seeds as our gold standard, we aligned the FLPSSs to PfamSDB.

Our data shows that 78% of the hits were TPs. The precision-recall curve of Foldseek probability showed that the recall (or sensitivity) changes less than 0.001 as we adjust the probability thresholds, while the precision changes from 0.78 to 0.82 when we change the Foldseek probability cutoff from its minimum (0.024) to its maximum (1). F-measure, the harmonic mean of Precision and Recall, provides a balanced assessment of a classification model’s performance. The Foldseek probability of 1 resulted in the highest F-measure which is 0.89.

Foldseek probability is a mapping between the alignment bitscore and the probability that the query and target belong to the same SCOPe superfamily (Personal communication with Milot Mirdita). We noticed that a Foldseek probability of 1 has been attributed to any alignment with a bitscore above 100. As the highest F-measure for Foldseek probability’s performance was achieved when Foldseek probability of 1 was considered, there is a chance that selecting a higher bitscore as the threshold would lead to a higher F-measure. We plotted the precision-recall curve by considering different bitscore thresholds rather than Foldseek probabilities, shown in Figure 2. The highest F-measure was 0.94 which was achieved by selecting the bitscore of 152 as the cutoff threshold. For the rest of the analysis, we only considered the Foldseek hits whose bitscore was greater than or equal to 152.

**Figure 2.**
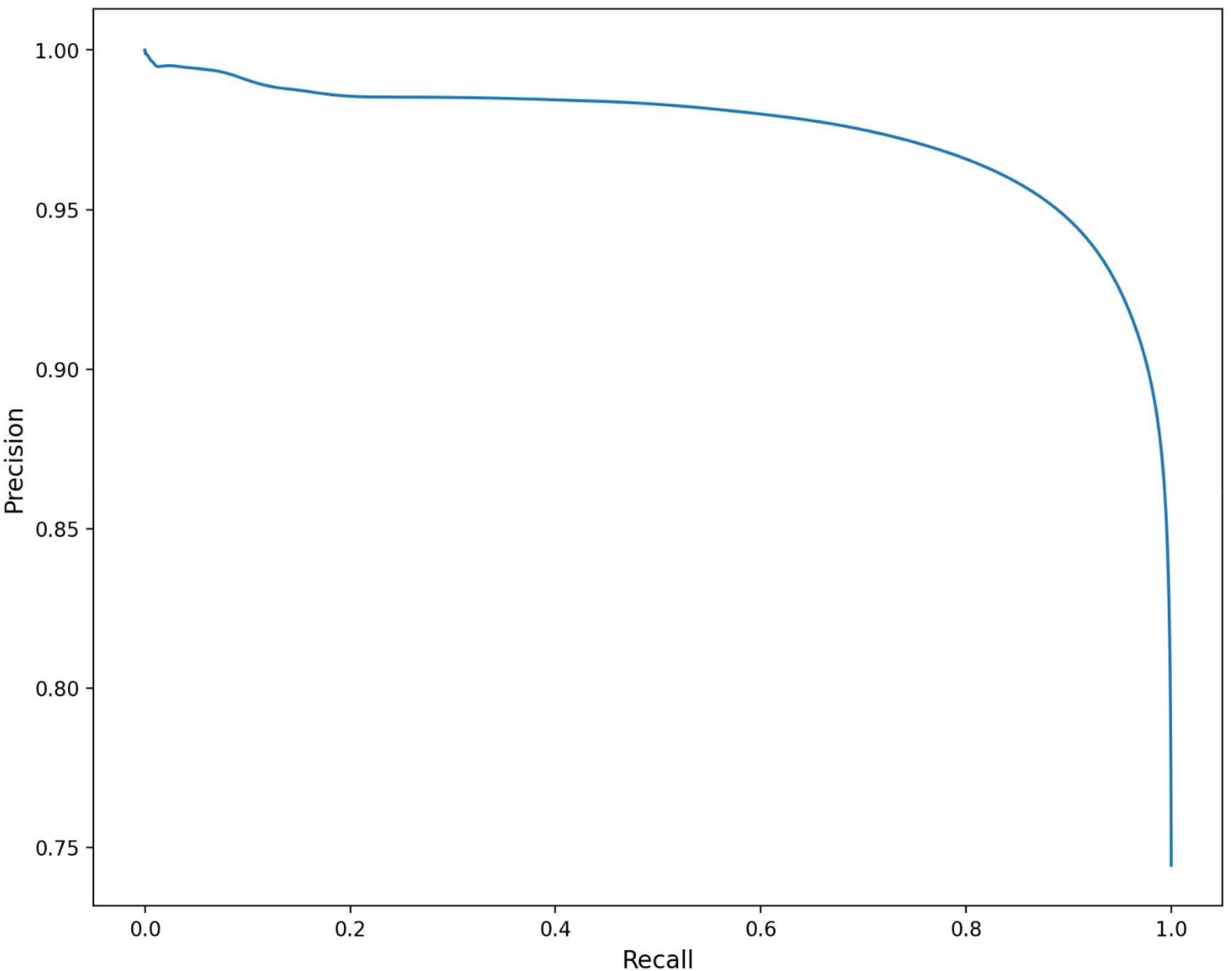
Precision-recall curve of Foldseek bitscore for differentiating TPs from FPs.

It is worth mentioning that here, we evaluated the precision and recall for all hits and found the optimum bitscore threshold based on FLPSS-against-PfamSDB alignment. There is also a single mapping between bitscore and Foldseek probability. However, the Pfam database considers different bitscore thresholds known as gathering thresholds for different Pfam domains, and the thresholds are manually curated. Although manual curation of the Foldseek bitscore threshold for each Pfam domain could enhance annotation performance, pursuing such an approach is beyond the purview of our current work.

As mentioned earlier, we also used MMseqs2, a sequence alignment tool, to see the added value of structural alignment. MMseqs2 does not report a probability score. Our benchmark showed that 98.9% of the alignments were TPs. The precision-recall curve showed the maximum F-measure (0.994) is achieved when the minimum bitscore (36) is used. So, we did not select any threshold on MMseqs2 hits.

### 3.3 Short or Low-Confidence Instances Less Likely to Identify Same Pfam Domain Matches

For each instance, the maximum number of TPs is equal to the number of instances in the same Pfam domain, a value we denote as N. Therefore, the maximum number of TPs for all instances of a particular Pfam domain amounts to *N*_2_. We then define the ‘proportion of retrieved instances’ as the ratio of the ‘total number of times instances of each Pfam domain were labeled as TP’ to *N* _2_. Figure 3 elucidates the relationship between this proportion of retrieved instances and two key parameters: the Average Length and the Average of “avg_pLDDT of instances”, per Pfam domain. According to Figure 3, Pfam domains with longer average lengths and higher avg_pLDDT generally display a higher propensity to retrieve all instances of the same Pfam domain from the query. This observation is in line with the findings of (Monzon, Paysan-Lafosse et al. 2022), where short proteins and proteins rich in residues with low pLDDTs exhibited a reduced likelihood of identifying their best reciprocal hits when examining the reciprocal best structural hits across two organisms’ proteomes.

**Figure 3.**
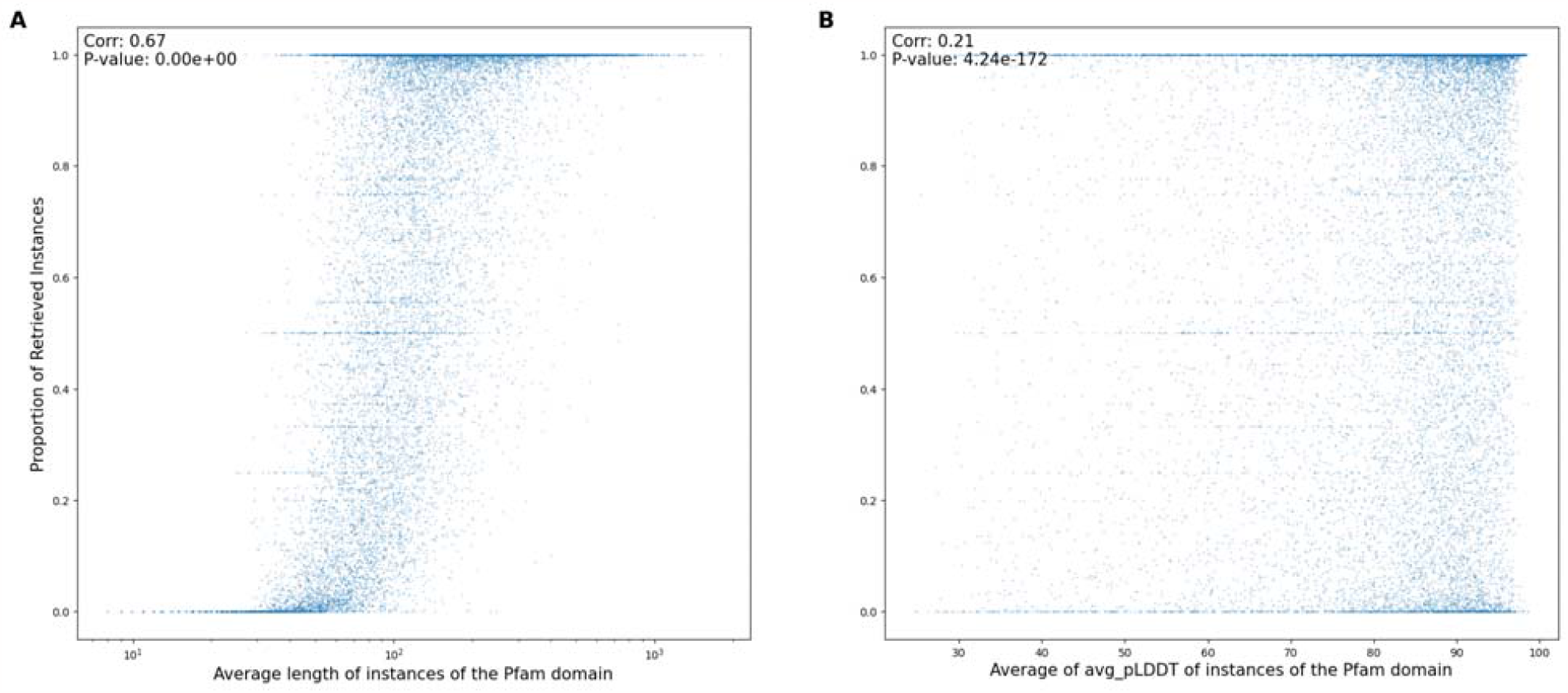
A: Relationship between the proportion of retrieved instances and the average length of instances for each Pfam domain. B: Relationship between the proportion of retrieved instances and the average avg_pLDDT of instances for each Pfam domain. The Spearman correlation coefficient, depicted in the upper left corner of each subfigure, provides a measure of the strength and direction of the relationship between the axes of each plot.

In certain Pfam domains, instances, despite having a substantial length (>100) and a high avg_pLDDT (>80), failed to retrieve any matches within the same Pfam domain - not even themselves. These particular instances predominantly exhibited low-complexity structures characterized by a profusion of helices. Further analysis revealed that the 3Di transformations of these structures also manifested this low complexity; most were sequences representing a single 3Di state. The 3Di structural alphabet, utilized by Foldseek, provides a representation of tertiary interactions between residues in a protein’s spatial configuration. This offers enhanced information density and fewer false positives than traditional backbone structural alphabets (van Kempen, Kim et al. 2023). During prefiltering, Foldseek identifies matches with 3Di sequences similar to the query protein using sequence alignment. Notably, Foldseek’s logs indicate that it automatically masks low-complexity regions during the 3Di state prefiltering. This likely explains the inability of low-complexity regions to align with related matches.

### 3.3 Domain Prediction: Comparable in Count to Pfam v35

We aligned the *T. brucei* proteins with the Pfam instances database, selecting non-overlapping hits as domain annotations. Figure 4 illustrates the number of domains annotated using various approaches according to which the number of predicted domains by Foldseek is shown to be comparable to that of Pfam v35 and is 25% more than the MMseqs2 predictions.

**Figure 4.**
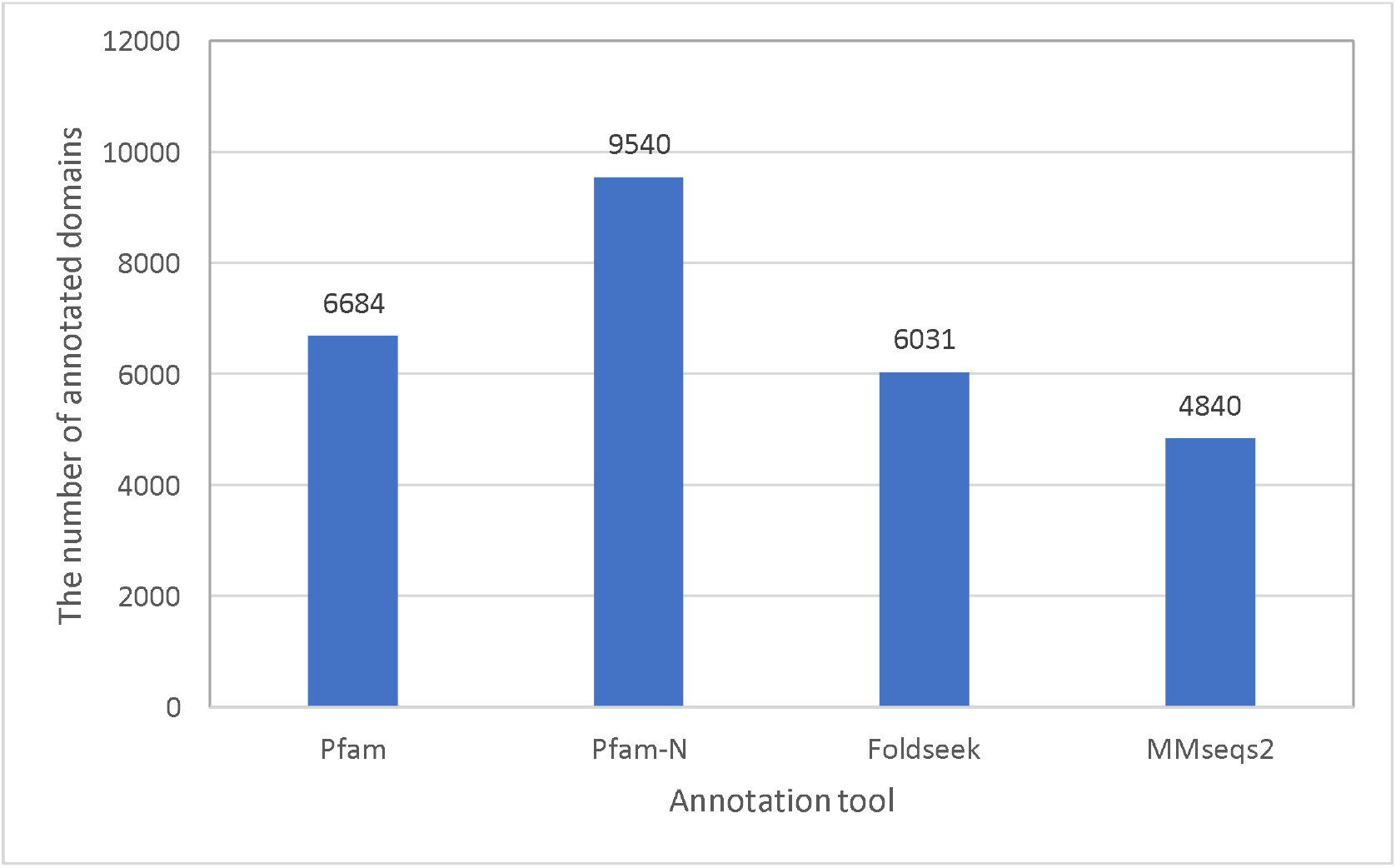
The number of domains annotated in different approaches.

Figure 5 presents precision and recall at both Pfam and Clan levels, using either Pfam or Pfam-N as the gold standard. The Pfam database groups different Pfam domains with similar sequence signatures into the same Clan and Clan-level statistics are also depicted in Figure 5. Compared to MMseqs2 annotations, those from Foldseek exhibit higher recall but lower precision. In all scenarios, Clan level precision exceeds 90%. This suggests that even when a Pfam domain identical to the gold standard may not be predicted based on structure, a quite similar Pfam domain is often attributed to the same region.

**Figure 5.**
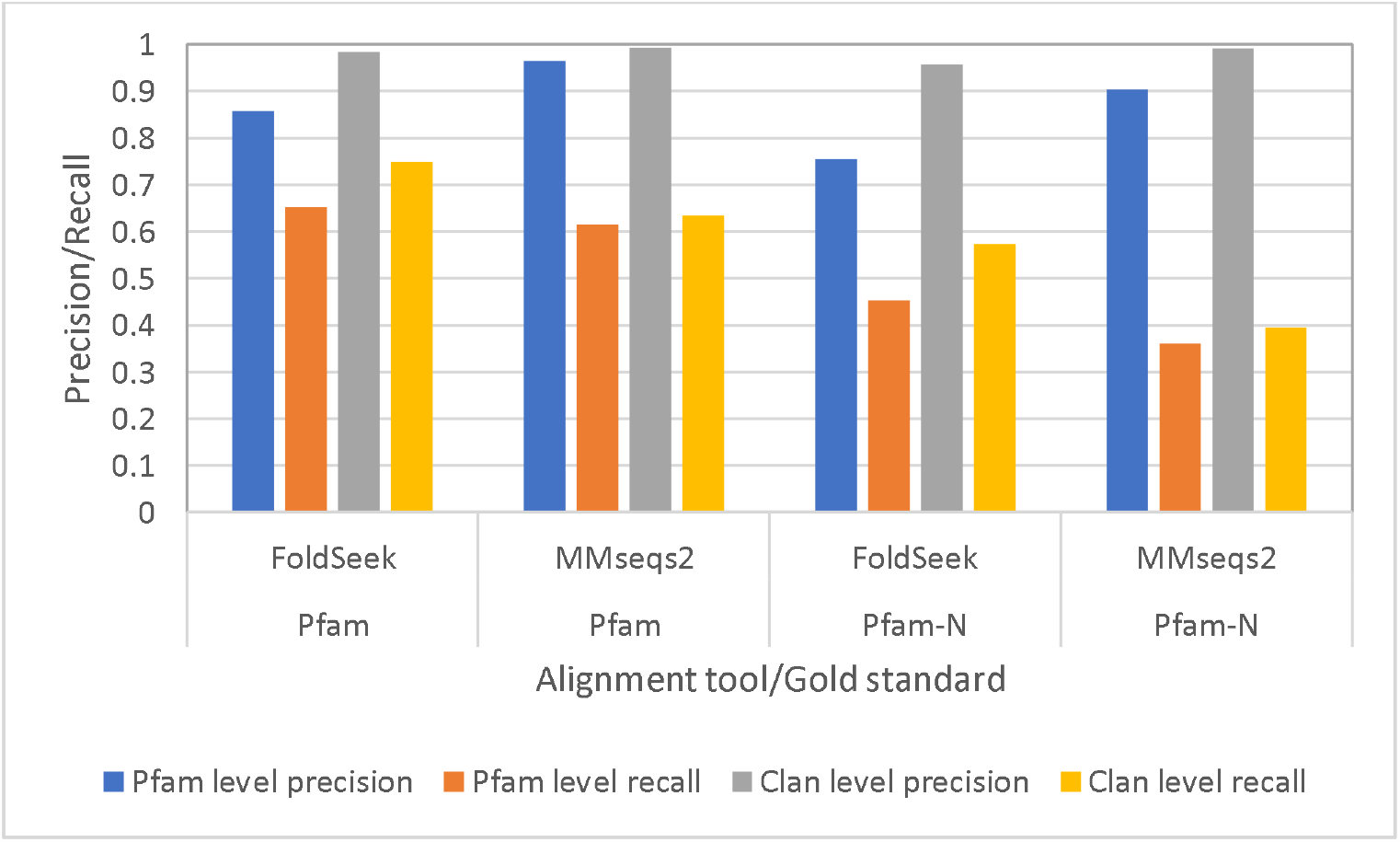
Precision and recall of domain annotations by MMseqs2 and Foldseek, using either Pfam or Pfam-N as the gold standard. In each panel, the top line indicates the annotation method, while the bottom line denotes the gold standard. For this figure, domain annotations corresponding to the regions utilized as Pfam seeds were excluded.

Figure 5 demonstrates that when using Foldseek for domain annotation, recall improved by just under 4% in comparison to MMseqs2. Yet, previous studies have shown that Foldseek substantially outperforms BLASTP, a comparable sequence-based tool. For instance, Monzon et al. employed both Foldseek and BLASTP to determine the best reciprocal relationship among multiple model organisms (Monzon, Paysan-Lafosse et al. 2022). Notably, since Foldseek utilizes MMseqs2 in its high sensitivity mode, it is crucial for a balanced comparison of sequence and structure-based alignment to also run MMseqs2 in this mode. Endeavoring to provide such a comparison, we examined the best reciprocal relationships among the organisms featured in Monzon et al.’s study. Our results indicate that by using MMseqs2 in its high sensitivity setting (-s 8.5), the best reciprocal relationship for an additional 200 proteins can be identified, specifically when analyzing the proteomes of *Drosophila melanogaster and Homo Sapiens*. Despite this, the majority of statistics from the Monzon et al. study remained consistent with our experiment. Furthermore, while running MMseqs2 in high-sensitivity mode did yield more “best reciprocal matches, Foldseek still significantly outstripped MMseqs2 in terms of established relationships. This implies that the modest edge Foldseek has over MMseqs2 in domain annotation might be influenced by the distinct biological questions under investigation.

### 3.4 Domain length impacts the recall significantly

**Error! Reference source not found**. Figure 6 illustrates the analysis of Pfam domains using Foldseek, MMseqs2, and HMMER. In Figure 6A, domains located in areas with low pLDDT scores show reduced detection by Foldseek, indicated by a lower recall rate. Notably, MMseqs2 (Figure 6C) also displays low recall but high precision for domains in low pLDDT regions. This can be attributed to the fact that IDR regions exhibit low sequence similarity (resulting in low recall), but sequences resembling an IDR are indeed IDRs (resulting in high precision).

**Figure 6.**
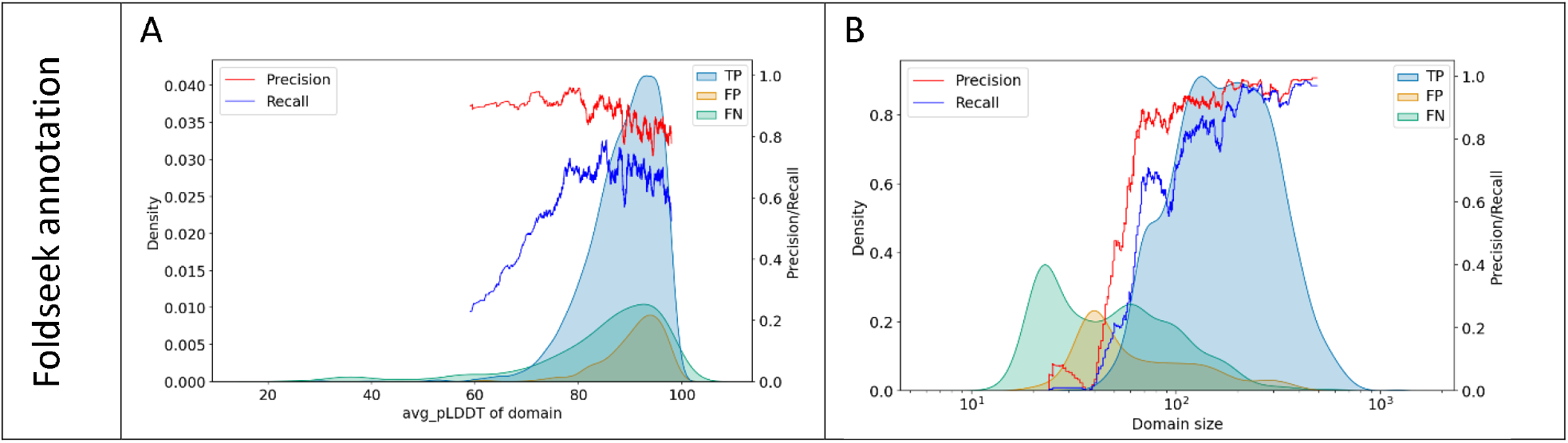
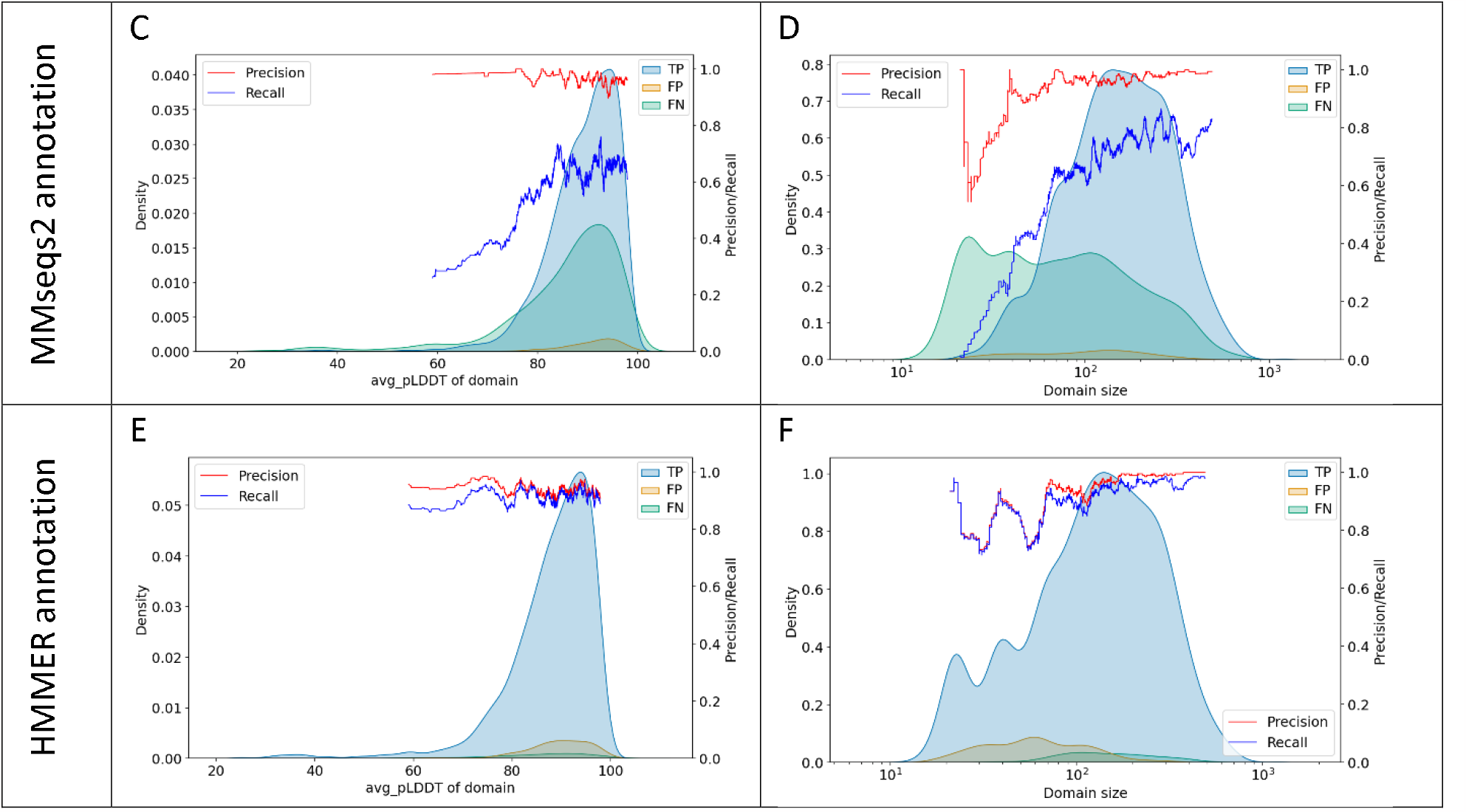
Analysis of Pfam Domains: Foldseek, MMseqs2, and HMMER Annotations. A: avg_pLDDT distribution for Foldseek domains. B: Size distribution of Foldseek domains. C: avg_pLDDT distribution for MMseqs2 domains. D: Size distribution for MMseqs2 domains. E: avg_pLDDT distribution for HMMER domains. F: Size distribution for HMMER domains. Distributions are colored according to their relationship with the gold standard (True Positive (TP), False Positive (FP), False Negative (FN)). Precision and recall are calculated using a rolling method for every 200 consecutive hits.

Figure 6B further demonstrates that the same observation holds true for the “domain size,” with small domains having a notably low recall rate. A direct comparison between Figure 6B and Figure 6D reveals that Foldseek does not enhance recall when retrieving small-sized domains, relative to MMseqs2; in fact, precision is even lower with Foldseek. This phenomenon can be attributed to the tendency of short domains not to fold into unique structures. Consequently, their structural resemblance to instances of different Pfam domains may increase the chance of alignment with a non-corresponding Pfam domain instance, thereby reducing precision. The same description can explain why the comparison of Foldseek and BLASTP for finding the best reciprocal hits between two organisms shows that many best reciprocal hits that are exclusively found by BLASTP, are less than 200 amino acids long (Monzon, Paysan-Lafosse et al. 2022).

Alignment bitscore depends on the length of the alignment and we expect that short domains would align with a lower bitscore even by HMMER, the program used by Pfam for domain annotations. However, since Pfam uses domain-specific bitscore thresholds, this can aid in annotating short domains. Indeed, Figure 6F shows that the HMMER top hit selection without considering the gathering threshold results in lower Precision for shorter domains. However, the reduction is less significant than Foldseek and MMseqs2.

Supplementary Figure 1 supports these observations, showing that similar patterns persist when Pfam-N is considered the gold standard. Additionally, Supplementary Figure 1 indicates that when Pfam-N is the gold standard, the precision of predictions for domains located in regions with low avg_pLDDT is lower than that of domains in high avg_pLDDT regions. The same figure also shows that the HMMER top hit selection has low precision and recall for retrieving short Pfam-N domains.

The Spearman correlation coefficient between avg_pLDDT and domain size for Pfam and Pfam-N domains is -0.076 and 0.79 respectively, showing that there is no meaningful relationship between them.

While Figure **6Error! Reference source not found**.B shows that the precision and recall for annotating long domains is relatively high compared to short domains, Figure 7A shows that long domains attributed by Foldseek have a slightly lower precision than shorter ones. This discrepancy may arise because long Foldseek domains, labeled as FP, often correspond to regions where shorter Pfam domains are typically predicted, a relationship further illustrated by Figure 7B. Supplementary Figure 2 shows similar patterns by considering Pfam-N as the gold standard. For example, Pfam-N predicts PF01909, Nucleotidyltransferase domain, and PF03828, Cid1 family poly A polymerase, for the N and C-terminal of Q38CM2, respectively. On the other hand, Foldseek predicts PF04928, Poly(A) polymerase central domain, for a region encompassing both regions.

**Figure 7.**
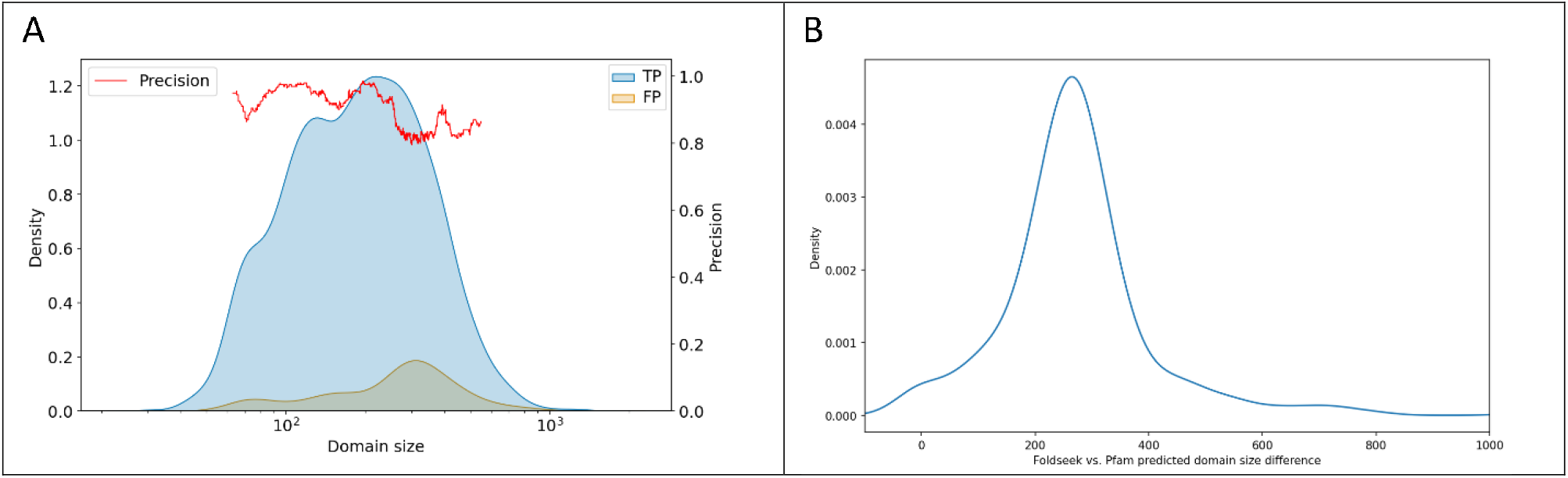
Analysis of Foldseek domains by Pfam. A: The distribution of length of Foldseek domains categorized by their relationships to the Pfam annotations (True Positives (TP), False Positive (FP)). B: The difference between the size of the false positive Foldseek domains and the Pfam domains predicted for the same region. Foldseek domains longer than 200 amino acids have been used for part B.

To assess the impact of choosing a bitscore threshold on domain annotation, we evaluated the precision and recall without imposing any threshold. Using Pfam as the gold standard, the lack of a bitscore condition increased recall by 0.8% but decreased precision by 2.4%. Similarly, when using Pfam-N as the gold standard, recall increased by 1.4%, while precision dropped by 3.4%. These findings suggest that employing the optimum bitscore threshold, as calculated from FLPSS-against-PfamSDB alignments, can enhance *T. brucei* domain annotation.

### 3.5 By increasing the e-value threshold, the majority of domains align with at least one related Pfam domain instance

As the next step, we explored if by using less stringent e-value thresholds, more domains could be annotated. Figure 8 illustrates how adopting less stringent e-value cutoffs in Foldseek leads to more domains aligning with at least one related seed. While this trend is also observed in MMseqs2 hits, the rate of increase relative to the e-value threshold is less pronounced than in Foldseek. However, using less stringent e-values may result in random structures aligning with query proteins, thereby raising the likelihood of inaccurate matches. Consistently, our benchmarking revealed that relying on hits with higher e-values leads to low-precision domain annotations, as anticipated (data not shown).

**Figure 8.**
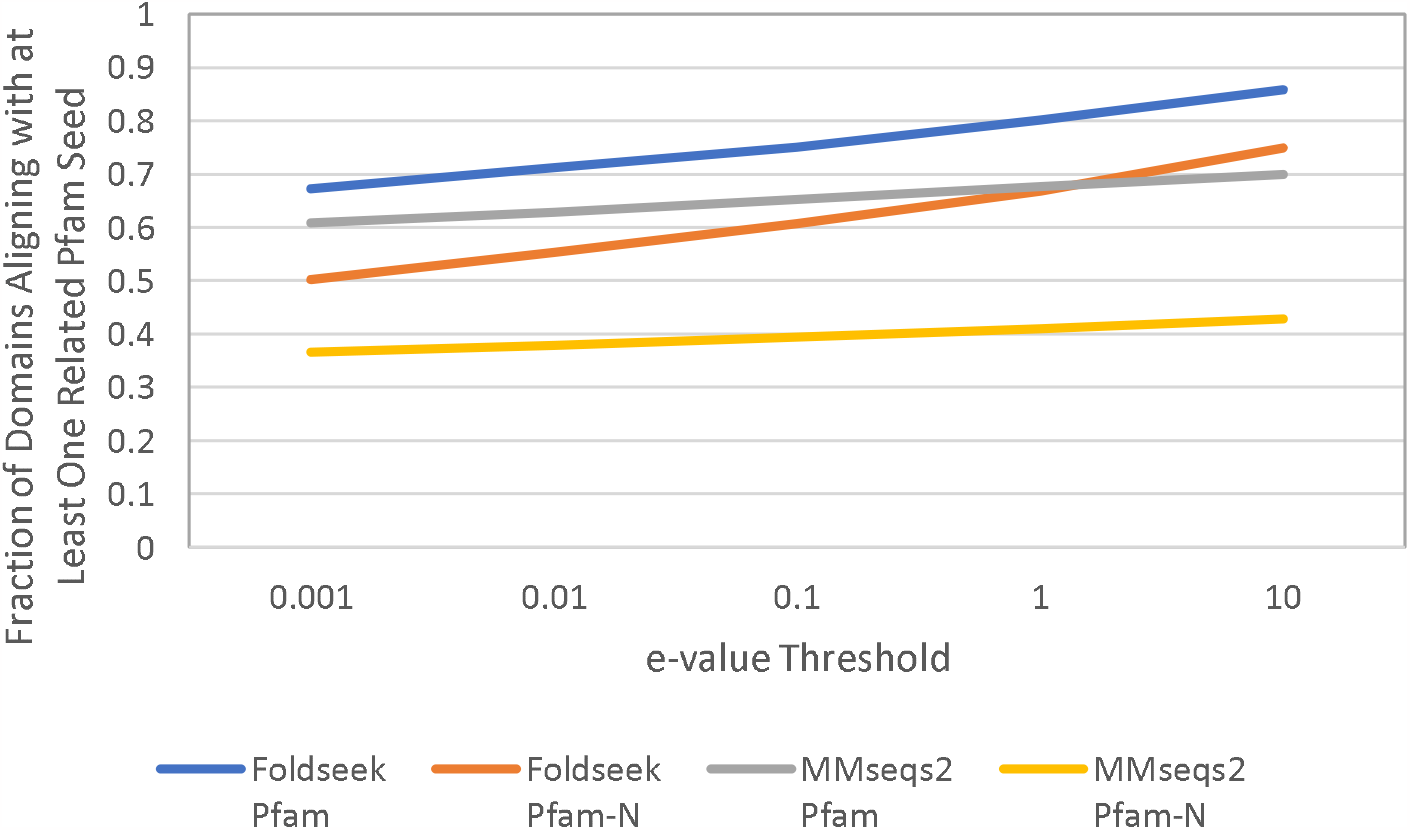
Comparison of the proportion of domains aligning with at least one related Pfam domain instance, using either Pfam or Pfam-N as the gold standard. The top line in the legend indicates the alignment tool used for domain annotation, while the bottom line specifies the gold standard.

Our findings underscore a complex challenge in structure-based domain annotation. Specifically, while using more lenient e-values often results in the query structure aligning with instances of the same Pfam domain, the aligned seed does not necessarily rank higher than those of other structurally similar domains.

In an attempt to improve the ranking of the hits, after aligning the FLSs with the domain database domain, we tried training an Artificial Neural Network (ANN) to predict if the query and target have the same Pfam domain using alignment characteristics such as sequence identity, LDDT, and bitscore as input features. However, rescoring using the probability of trained ANN did not improve the precision significantly (data not shown). We speculate that re-ranking based on the conservation of critical residues could potentially enhance precision in future attempts.

Observing a sharp increase in domains that structurally align with at least one related hit— contrasting with MMseqs2 domains—we hypothesized that the e-value of hits would generally be higher when a structural alignment is performed, compared to sequence alignment. Figure 9, illustrating the log of the ratio of e-values for identical “query and target” pairs, supports this hypothesis: in 78% of cases, the Foldseek e-value exceeds the MMseqs2 e-value. This phenomenon can be attributed to Foldseek’s scoring system, which takes into account both structural and sequence similarity. Since the complexity of possible protein folds in the universe is lower than that of sequences (Govindarajan, Recabarren et al. 1999), we expect the query protein to align with a larger number of proteins in the database based on structural similarity. Consequently, for each query structure, there is an increased likelihood of obtaining random hits, effectively leading to less significant e-values.

**Figure 9.**
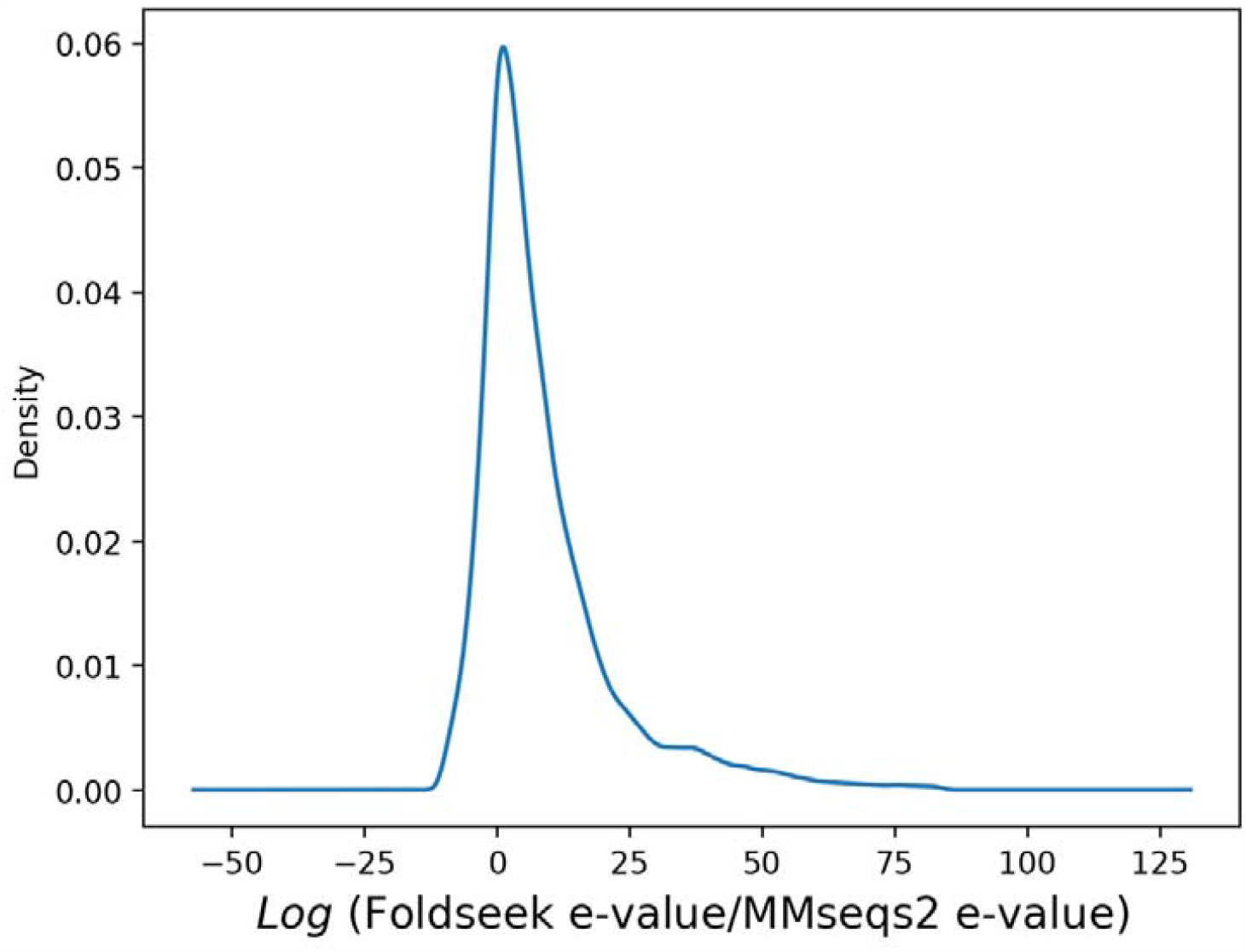
The ratio of Foldseek e-value to the MMseqs2 e-value for identical “query and target” pairs. the top and bottom 1% of data points have been excluded to highlight the overall distribution.

### 3.6 New domains predicted above domains predicted by Pfam and Pfam-N

In regions that lacked Pfam predictions, MMseqs2 pinpointed 265 domains. Similarly, it found 331 domains where Pfam-N had not identified any. Furthermore, in areas untouched by both Pfam and Pfam-N, MMseqs2 unveiled 134 new domains with an average length of 40.3. An examination of the MMseqs2 annotations reveals that the average sequence identity for the domains it annotated is 80.5%. In 54 of these cases, the query and target are 100% identical. This means that, in some instances, even when part of the query sequence has been utilized as a Pfam seed, the corresponding domain remains undetectable in the query protein by both Pfam and Pfam-N. For example, position 469 to 497 of the protein Q9N937 has been used as a Pfam seed for the domain PF00560 (Leucine Rich Repeat). However, Pfam has not annotated the same region probably because they had not satisfied the gathering thresholds.

Foldseek identified 1113 domains in areas where Pfam made no predictions and 523 domains in areas without Pfam-N predictions. Additionally, Foldseek detected 403 new domains in regions where both Pfam and Pfam-N had not previously identified any domains. These domains are notably longer, with an average length of 254.9 amino acids. Additionally, 177 of the proteins annotated by Foldseek in TriTrypDB (Amos, Aurrecoechea et al. 2022) are described as “hypothetical protein, conserved.” The most frequently attributed domain is PF13458, known as the “Periplasmic binding protein.” PF13458 has been attributed 32 times. According to the TriTrypDB website, this particular Pfam domain is primarily associated with genes described as “Adenylate cyclase.” Interestingly, in our workflow, all the genes that were annotated with PF13458 also included “Adenylate cyclase” in their description.

### 3.7 Case study: Domains predicted for proteins involved in *T. brucei* mtRNA editing

Trypanosomes possess a unique mitochondrial DNA structure composed of multi-kilobase-size fragments. These fragments are referred to as maxicircles, which are interconnected with numerous smaller minicircles (Aphasizheva, Alfonzo et al. 2020). Within the maxicircles, 12 of the encoded genes are classified as cryptogenes, signifying that they cannot be directly translated after transcription. During the editing process, multiple uracil (U) bases must be either added or deleted to render these genes ready for translation (Aphasizheva, Alfonzo et al. 2020). This intricate editing process is guided by guide-RNAs (gRNAs), which are primarily encoded by the minicircles. The gRNAs operate by marking the editing sites on the substrate RNA. They anneal to the substrate, creating bulges that signal the specific locations for catalytic enzymes to modify. For a comprehensive review of the editing process please refer to (Aphasizheva, Alfonzo et al. 2020). As a case study, we look deeper into the domains predicted for some of the proteins involved in mtRNA editing.

The prediction suggests that KREPA4 (UniProt ID: Q38B91) may contain the PF00436 domain, known as “Single-strand binding domains (SSB).” Pfam has also predicted PF00436 for KREPA6 (UniProt ID: Q38B90), with this prediction being consistent with that of Pfam-N. KREPA4 and KREPA6 are adjacent in the genome, and they appear as the closest non-self structural hits to each other when searched against the structure of all organism proteins. This suggests the possibility that they might be paralogs. The expectation that paralogous proteins contain similar domains supports this case. Analysis of the sequence signature of PF00436 reveals that positions 8, 69, and 76 are conserved and contain the amino acid Glycine. Correspondingly, these positions in KREPA4 also contain Glycine. The consistency in this specific amino acid placement between KREPA4 and the conserved signature of PF00436 provides evidence to support the prediction of this domain.

Both KREX1 (UniProt ID: Q57WU3) and KREX2 (UniProt ID: Q38BP2) are predicted to contain PF03159, which corresponds to the XRN 5’-3’ exonuclease N-terminus domain. This prediction is consistent with that made by Pfam-N for these proteins. An early study had hypothesized a similar function for these proteins (referred to as MP100 and MP99 in the publication), and this hypothesis was based on the conservation of certain amino acids. However, the confidence in attributing this domain to KREX1 and KREX2 had been limited at the time due to the low overall homology with the known XRN domains (Worthey, Schnaufer et al. 2003).

We predicted the presence of PF02940, identified as the “mRNA capping enzyme, beta chain,” in RESC1 (UniProt ID: Q57XL7) and RESC2 (UniProt ID: Q586X1). The beta chain of the mRNA capping enzyme is known for its triphosphatase activity. A recent study by Dolce et al. elucidated the structure of the RESC1-2 complex using cryo-electron microscopy (Dolce, Nesterenko et al. 2023). The researchers reported a structural similarity between RESC1-2 and RNA capping enzymes, which typically have cationic cofactors and are involved in reactions that release phosphate. The study further examined the charged residues within the tunnels of the RNA capping enzyme, noting that one-half of the pattern interacts with the cationic cofactors, and the other half with the released phosphate. However, this pattern was not observed in RESC1-2, leading to the assessment that RESC1 and RESC2 are unlikely to be active enzymes (Dolce, Nesterenko et al. 2023).

For RESC5 (UniProt ID: Q389F5), the prediction includes PF19420, identified as “N,N-dimethylarginine dimethylhydrolase (DDAH) within eukaryotes”, a domain related to arginine metabolism. This prediction was bolstered by a recent study that succeeded in crystallizing the structure of RESC5, revealing its structural similarity to the DDAH fold (Salinas, Cannistraci et al. 2023). However, this same study also uncovered key differences. Most notably, RESC5 was found to lack residue conservation in critical positions that are otherwise characteristic of the DDAH fold. Further investigation into RESC5’s interaction with the DDAH substrate and product provided additional insights. The researchers conducted a Thermal shift assay, a technique used to assess protein-ligand interactions. The addition of the DDAH substrate and product to RESC5 had no discernible effect on the assay’s results. This lack of effect serves as an indicator that there is no interaction between RESC5 and the DDAH substrate or product (Salinas, Cannistraci et al. 2023). The findings from this detailed analysis demonstrate that despite superficial similarities, RESC5 likely does not function in the same manner as DDAH.

## 4 Conclusion and future work

In this study, we developed a database of domain structures by segmenting the structures predicted by Alphafold at their domain boundaries. To annotate domains, we structurally aligned query proteins with this domain database using Foldseek. Subsequently, we selected the highest-scoring, non-overlapping hits. We then benchmarked these predictions against Pfam v35.0 and Pfam-N predictions.

Our data indicates that for short domains (those less than 100 amino acids in length), structure-based domain annotation is imprecise. This lack of precision can be attributed to the fact that short sequences often lack distinctive folds. Consequently, there is a heightened likelihood of random structural similarities between different domains, resulting in reduced precision. Given that a significant fraction of domains are short, our results suggest that while structure-based Pfam annotation cannot supplant sequence-based domain annotation, it can complement it, particularly when annotating longer domains. We are keen to explore the synergy between sequence-based and structure-based domain annotations in future studies.

From an organism-specific standpoint, our study offers insights into the potential functions of genes in *T. brucei*. These insights are ripe for experimental validation. One standout prediction is the anticipated 5’-3’ exonuclease activity for KREX1 and KREX2; we are eager to corroborate this finding through wet lab experiments.

## 5 Data availability

The scripts for creating the PfamSDB are available at https://github.com/Pooryamb/MakingPfamSDB. Scripts for benchmarking can be found at https://github.com/Pooryamb/BenchmarkingFS/.

## 6. Acknowledgments

We extend our heartfelt gratitude to the reviewers for their insightful comments and constructive feedback, which significantly contributed to enhancing the quality of this manuscript. Special thanks go to Dr. Naghmeh Poorinmohammad for her invaluable comments. This project has been supported by a research grant 252733 from CIHR to RS. Robert Harpur Foundation is also acknowledged for providing financial support to PMB.

## Notes

https://github.com/Pooryamb/MakingPfamSDB

https://github.com/Pooryamb/BenchmarkingFS

### Competing Interest Statement

The authors have declared no competing interest.

### Summary of Updates

In the first version of the manuscript, we trained a Random Forest to distinguish True Positives from False Positives based on their alignment characteristics. Domains annotated by the Pfam for Trypanosoma brucei were used for this training. Upon revisiting our approach during revisions, we found that we could achieve superior performance by setting an optimal bitscore threshold. To determine this optimal threshold, we aligned full-length proteins from Pfam seed sources against the Pfam seeds, aiming for an organism-unspecific threshold to differentiate True Positives from False Positives. We enhanced the benchmarking of our domain prediction method, comparing it more rigorously against established domain annotation tools, namely Pfam v35.0 and Pfam-N. Additionally, we comprehensively compare Foldseek with MMseqs2, its counterpart sequence aligner. Interestingly, while structural aligners are theoretically more sensitive than sequence aligners, we found that domain annotation by Foldseek trails the predictions made by sequence-based domain annotation tools such as HMMER (used for predicting Pfam domains) and ProtENN (used for predicting Pfam-N domains) in Trypanosoma brucei, and we have explained the reasons for this observation.

