## Supplementary Figure 1 for "Functional domain annotation by structural similarity"

| Foldseek annotation | A  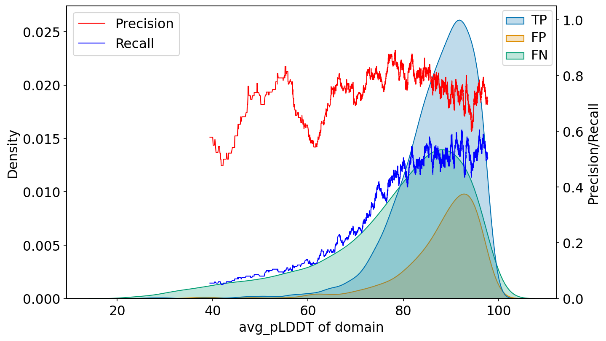 | B  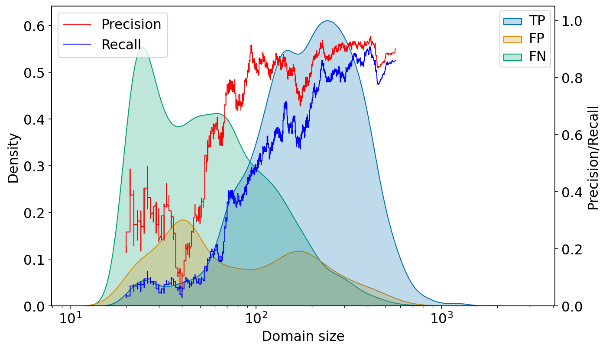 |
| --- | --- | --- |
| MMseqs2 annotation | C  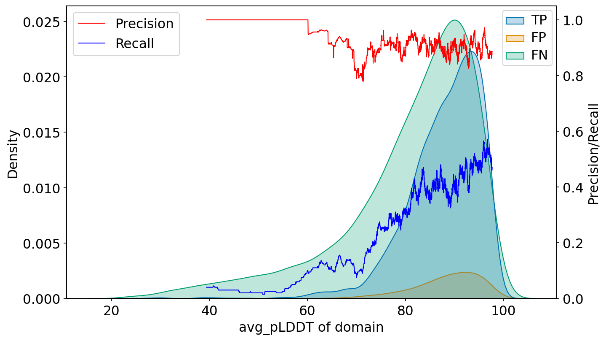 | D  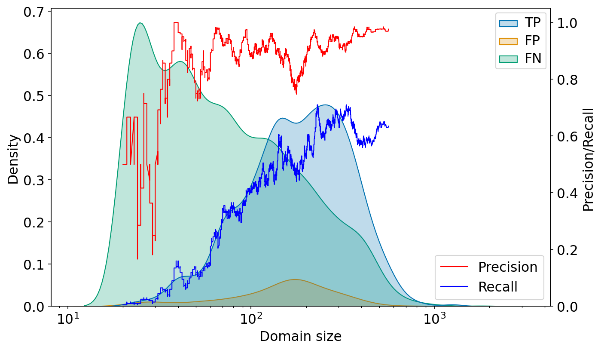 |
| HMMER annotation | E  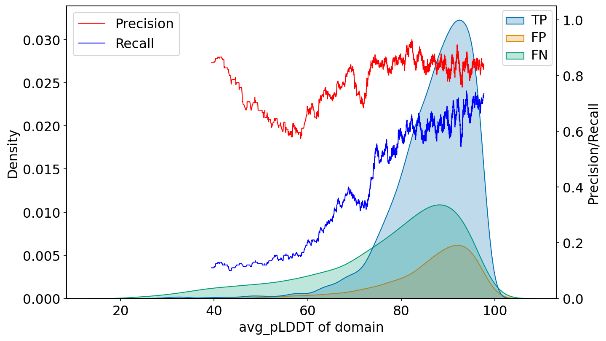 | F  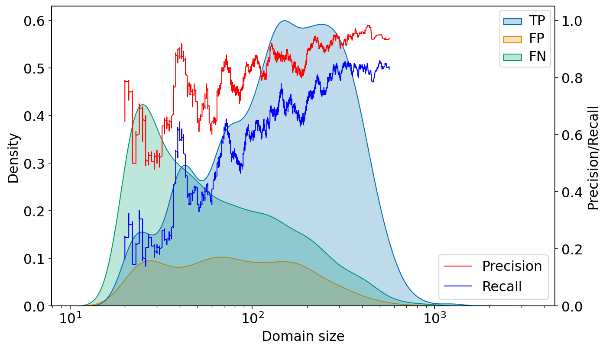 |

Supplementary Figure 1. Analysis of Pfam-N Domains: Foldseek, MMseqs2, and HMMER Annotations.

A: avg_pLDDT distribution for Foldseek domains (TP, FP, FN). B: Size distribution of Foldseek domains.

C: avg_pLDDT distribution for MMseqs2 domains. D: Size distribution for MMseqs2 domains.

E: avg_pLDDT distribution for HMMER domains. F: Size distribution for HMMER domains. Distributions are colored according to their relationship with the gold standard (True Positive (TP), False Positive (FP), False Negative (FN)).

Precision and recall are calculated using a rolling method for every 200 consecutive hits.
