## Supplementary Figure 2 for "Functional domain annotation by structural similarity"

| A  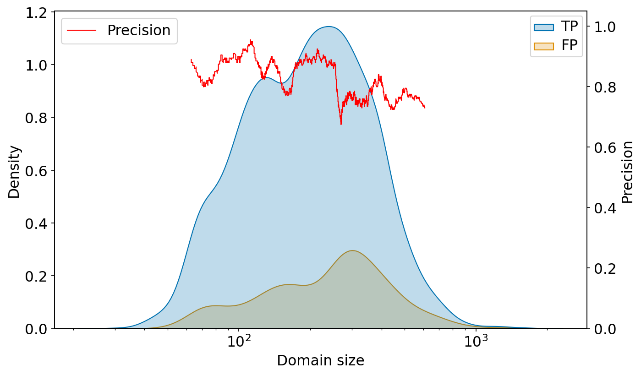 | B  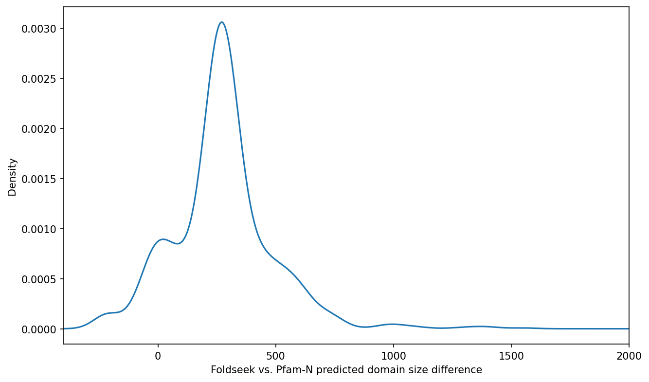 |
| --- | --- |

Supplementary Figure 2. Analysis of Foldseek domains by Pfam-N. A: The distribution of length of Foldseek domains categorized by the Pfam-N annotations (True Positives (TP), False Positive (FP)). B: The difference between the size of the false positive Foldseek domains and the Pfam-N domains predicted for the same region. Foldseek domains longer than 200 amino acids have been used for part B.
